# Expansion and Differentiation of Adult Human Pancreas-Derived Progenitor Cells into Functional Islet-Like Organoids

**DOI:** 10.64898/2026.04.14.718604

**Authors:** Jayachandra Kuncha, Carly M. Darden, Jeffrey T. Kirkland, Jean-Philippe Blanck, Kelli Fowlds, Michael Cho, Juan S. Danobeitia, Bashoo Naziruddin, Michael C. Lawrence

**Affiliations:** Islet Cell Laboratory, Baylor Scott & White Research Institute, Dallas, TX; Annette C. and Harold C. Simmons Transplant Institute, Baylor University Medical Center, Dallas, TX; Baylor Institute for Immunology Research, Dallas, TX; Department of Bioengineering, University of Texas at Arlington, Arlington, TX

## Abstract

**Background and Aims:** Adult pancreas-derived islet progenitor cells (IPCs) have recently been shown to expand in culture and differentiate into endocrine-like organoids. However, translation of this approach to a clinically compatible workflow requires cell enrichment strategies and validation using tissue obtained during real-world clinical procedures. Here, we adapted our previously described IPC platform to non-endocrine pancreatic tissue fractions generated during clinical islet isolation procedures and evaluated their capacity to generate functional islet organoids.

**Methods:** Non-endocrine pancreatic tissue fractions obtained during clinical islet isolation were expanded ex vivo and enriched using fluorescence-activated cell sorting (FACS) for CD81 and CD9, surface markers previously identified in IPC populations. Sorted cells were expanded, induced to form IPC clusters, and differentiated with ISX9 to generate islet organoids. Differentiation was assessed by gene expression analysis, flow cytometry, immunofluorescence, calcium flux assays, glucose-stimulated insulin and glucagon secretion, and single-cell RNA sequencing.

**Results:** Clinically derived non-endocrine cell fractions yielded expandable IPC populations expressing progenitor-associated markers. FACS-purified and expanded CD81^+^/CD9^+^ IPCs were enriched with BMPR1A and P2RY1. Sorted cells generated three-dimensional BMPR1A^+^ and RGS16^+^ IPC clusters. IPC clusters differentiated into islet organoids with upregulated expression of canonical beta-and alpha-cell transcription factors. Single-cell transcriptomic profiling revealed activation of coordinated endocrine gene programs and alignment with reference human islet endocrine signatures, while the undifferentiated IPC compartment was marked by enrichment of PTX3, FST, CEMIP, and GREM1. Terminally differentiated cells exhibited depolarization-induced calcium influx and glucose-regulated insulin and glucagon secretion.

**Conclusions:** These findings establish an adaptable workflow for expansion and production of functional islet organoids recovered from clinically derived pancreatic tissue. This strategy may provide an unlimited autologous source of adult progenitor-derived islets for future islet cell replacement therapies in diabetes.

## INTRODUCTION

Diabetes mellitus remains a major global health burden, and restoration of functional pancreatic islet mass is a central therapeutic objective. Clinical islet transplantation can re-establish glycemic control in selected patients (1, 2, Hering, 2025 #4180). However, donor scarcity, variability in islet yield, and limited long-term graft durability restrict broader application. In the setting of total pancreatectomy with islet autotransplantation (TPIAT), insufficient recovery of islet mass frequently results in persistent insulin dependence despite technically successful procedures (3). Strategies that enable expansion or regeneration of endocrine-competent cells from patient-derived pancreatic tissue may therefore improve clinical outcomes and broaden access to islet replacement therapies (4).

Recent advances in stem cell-derived islet platforms have demonstrated the feasibility of generating endocrine tissue ex vivo (5–8). In parallel, emerging evidence suggests that adult pancreatic tissue retains a degree of plasticity that can be harnessed under defined culture conditions (9–16). In our previous work, we identified an expandable population of adult pancreas-derived islet progenitor cells (IPCs) capable of forming three-dimensional clusters and differentiating into endocrine-like organoids (17). Transcriptomic profiling of these cells revealed features consistent with immature or dedifferentiated endocrine states, including enrichment of surface markers CD81 and CD9.

To translate this platform toward clinical application, it is necessary to identify progenitor-competent populations within pancreatic tissue obtained under real-world clinical conditions. During clinical islet isolation procedures for TPIAT, enzymatic digestion and purification generate non-islet tissue fractions that are not used for transplantation. We hypothesized that these clinically processed fractions contain cells capable of expansion and endocrine differentiation, and that CD81/CD9-based sorting could provide a practical strategy to enrich such populations.

In the present study, we adapted our IPC expansion and differentiation platform to non-endocrine pancreatic tissue fractions obtained during clinical islet isolation. We evaluated whether prospective enrichment using CD81 and CD9 yields populations capable of reproducibly generating IPC clusters and functional islet organoids. Differentiation was characterized using transcriptional, protein-level, and functional assays, including single-cell RNA sequencing, calcium flux analysis, and glucose-stimulated hormone secretion.

Collectively, these studies establish a clinically adaptable workflow for generating islet-organoids from patient-derived pancreatic tissue and provide a framework for future autologous cell expansion strategies in diabetes.

## MATERIALS AND METHODS

### Human subjects

The use of human subjects and associated procedures was approved by the Institutional Review Board of Baylor University Medical Center at Dallas. Total pancreatectomy with islet autotransplantation (TPIAT) was performed in patients meeting clinical criteria, including severe abdominal pain, chronic narcotic dependence, and impaired quality of life. Patients and legal guardians provided informed consent and were counseled regarding potential outcomes and complications associated with the procedure.

### Islet isolation and collection of non-endocrine pancreatic cell fractions

Pancreatic islet isolation was performed at the Baylor Scott & White Health (BSWH) cGTP Islet Cell Processing Laboratory according to established FDA guidelines. During the isolation process, non-endocrine pancreatic cell fractions were collected at multiple stages, including enzymatic digestion, enzyme dilution, tissue recombination, pellet washing, and COBE 2991 purification. Collected fractions were centrifuged at 2000 rpm for 5 minutes at 4°C to pellet dissociated cells. Cell pellets were resuspended in Dulbecco’s phosphate-buffered saline (DPBS; Ca²⁺/Mg²⁺; Thermo Fisher Scientific) and pooled for downstream processing. Islet-containing fractions were separately recombined and prepared for clinical infusion.

### Isolation and culture of dissociated pancreatic cells

Dissociated cells from non-endocrine pancreatic fractions were resuspended in RPMI supplemented with N-acetyl cysteine, Trolox, and retinoic acid (NTR) and 20% fetal bovine serum (FBS). Cell suspensions were filtered through a 40–60 mesh cell dissociation sieve (Sigma) to remove debris and fibrous material, followed by filtration through a 100 µm nylon filter (Corning) to remove cell aggregates. Filtering was repeated as needed until a single-cell suspension was confirmed by brightfield microscopy. Cells were centrifuged and resuspended in RPMI containing 10% FBS and seeded at a density of 9 × 10⁶ cells per 150 mm tissue culture dish. Cultures were maintained at 37°C with 5% CO₂ and ∼90% humidity.

### Expansion of pancreatic-derived cells

Following initial seeding, non-adherent cells were removed, and adherent cells were washed with phosphate-buffered saline (PBS) lacking calcium and magnesium. Fresh medium was added, and cells were cultured with medium changes every 48 hours. For passaging, cells were washed with PBS and dissociated using TrypLE Express (Gibco) for 7–10 minutes at 37°C. Cells were centrifuged, resuspended in RPMI with 10% FBS, counted, and reseeded into tissue culture-treated dishes. Cells were expanded for 2–3 passages under standard culture conditions.

### Flow cytometry analysis and cell sorting

Cells were dissociated into single-cell suspensions using TrypLE Express and filtered through a 70 µm nylon cell strainer. Cells were washed in FACS buffer (2% FBS in PBS), and viability was assessed using acridine orange/propidium iodide staining (Nexcelom Bioscience). For surface staining, 1 × 10⁶ cells were incubated with fluorophore-conjugated antibodies against CD81 and CD9. For intracellular staining, cells were fixed and permeabilized using Cytofix/Cytoperm (BD Biosciences) and stained for insulin and glucagon. Data were acquired using an LSR Fortessa flow cytometer (BD Biosciences) and analyzed using FlowJo software (v10.10.0). Fluorescence minus one (FMO) and unstained controls were used for gating. Cell sorting was performed using a BD FACSAria II to isolate CD81⁺/CD9⁺ and CD81⁻/CD9⁻ populations.

### Expansion of enriched IPC populations and formation of IPC clusters

Following sorting, cells were seeded onto Matrigel-coated culture dishes in RPMI containing 20% FBS and maintained under standard culture conditions for 48 hours without disturbance. Cells were passaged at ∼90% confluence and expanded for 2–3 passages, with medium changes every 48 hours. To generate IPC clusters, cells were cultured at high density in RPMI supplemented with NTR and 10% FBS for 14–21 days, with medium changes every 48 hours, resulting in formation of three-dimensional clusters.

### Isolation of IPC clusters

IPC clusters were separated from surrounding monolayer cells by gentle dissociation using diluted TrypLE Express (1:1 in PBS without calcium and magnesium) for 7–10 minutes at 37°C. IPC clusters were isolated using reversible cell strainers (Stemcell Technologies) with a 37 µm mesh size. IPC clusters were collected by reversing the strainer and flushing with complete medium.

### Differentiation of IPC clusters into islet cell organoids

IPC clusters were plated at approximately 500 clusters per 10 cm dish and cultured in RPMI supplemented with ISX9 (Cayman Chemical) and 2–5% FBS for 2–14 days to induce endocrine differentiation. Medium was replaced every 48 hours. Cell viability was assessed using fluorescein diacetate (FDA) and propidium iodide (PI) staining and calculated as the percentage of FDA-positive cells relative to total cells.

### Real-time quantitative PCR (RT-qPCR)

Total RNA was isolated using TRIzol (Life Technologies), and cDNA was synthesized from 1 µg RNA using a high-capacity cDNA reverse transcription kit (Thermo Fisher Scientific). Quantitative PCR was performed using a QuantStudio 7 Flex Real-Time PCR system (Applied Biosystems). Gene expression was calculated using the 2^−ΔΔCt method and analyzed in triplicate.

### Glucose-stimulated insulin and glucagon secretion (GSIS/GSGS)

Differentiated islet organoids were equilibrated in Krebs-Ringer bicarbonate HEPES buffer (KRBH) containing 1.67 mM glucose for 30 minutes at 37°C. Samples were then incubated in low (1.67 mM) or high (16.7 mM) glucose conditions for 1 hour. Insulin and glucagon secretion were measured using ELISA kits (Alpco). Total protein content was determined using a BCA assay, and secretion values were normalized to protein content and expressed as ng insulin/mg protein/hour and pg glucagon/mg protein/hour.

### Immunofluorescence

IPC islet organoids were differentiated on chamber slides and fixed in 4% paraformaldehyde for 10 minutes at 37°C. Cells were permeabilized using 0.1% Triton X-100 and blocked with 2% bovine serum albumin (BSA). IPC islet organoids were incubated overnight at 4°C with Alexa Fluor 488-conjugated anti-insulin antibody (Thermo Fisher Scientific). Samples were washed and mounted, and images were acquired using confocal fluorescence microscopy.

### Calcium flux analysis

Islet organoids were dissociated into single cells using TrypLE Express and filtered through a 70 µm strainer. Cells were resuspended in RPMI with 10% FBS and analyzed using an LSR Fortessa flow cytometer. Baseline calcium levels were recorded for 2 minutes, followed by depolarization with 50 mM KCl. Calcium responses were recorded for an additional 3 minutes. Data were analyzed using FlowJo in kinetic mode.

### Statistical analysis

Statistical analyses were performed using GraphPad Prism 10. Comparisons between groups were conducted using one-way ANOVA followed by Dunnett’s multiple comparisons test, as appropriate. Data are presented as mean ± standard deviation, and statistical significance was defined as p < 0.05.

## RESULTS

### Stepwise workflow for generation of IPC-derived islet organoids from non-endocrine pancreatic fractions

To establish a clinically adaptable platform for harvesting and expanding IPCs for generation of islet-like organoids, we applied a stepwise workflow to non-endocrine pancreatic cell fractions obtained during clinical islet isolation procedures (Fig 1). Harvested cell fractions were cultured for 2-3 weeks to enable expansion of IPCs (Fig 2A). Expanded cells were prospectively enriched by fluorescence-activated cell sorting (FACS) for CD81 and CD9. CD81^+^/CD9^+^ IPCs were then serially passaged to generate IPC monolayers, which gave rise to three-dimensional IPC clusters upon culture at high cell density (Fig 2B and C). Clusters were subsequently isolated from surrounding monolayer cells and advanced to endocrine differentiation conditions to generate functional islet-like organoids (Fig 2D). This workflow was designed to capture and recover a defined CD81^+^/CD9^+^ population associated with IPC-related features including expression of CD201 (PROCR), ALK3 (BMPR1A), and RGS16, as previously described (Darden et al.;Qadir et al; Wang et al). To further characterize these cultures, we examined the morphology and reproducibility of IPC expansion and cluster formation across independent samples.

**Figure 1.**
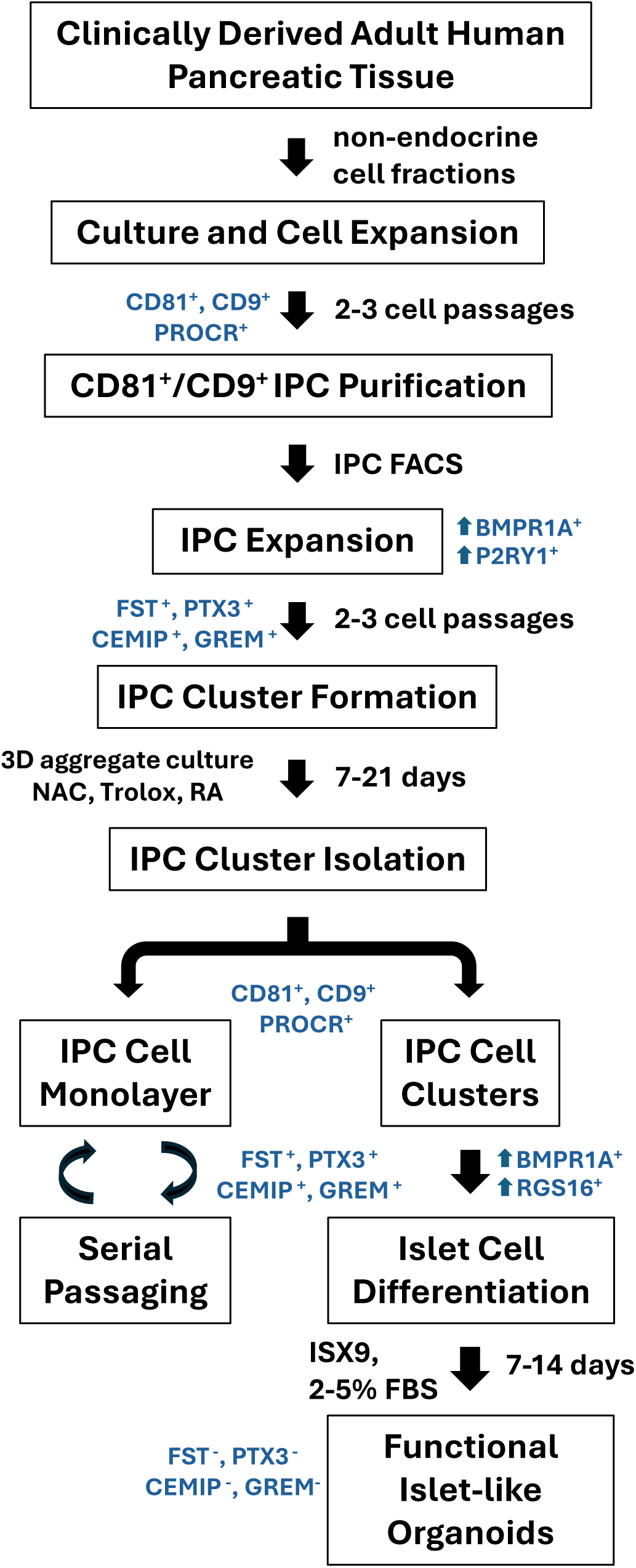
Workflow for generation of IPC-derived islet organoids from non-endocrine pancreatic fractions Schematic overview of the experimental workflow. Non-endocrine pancreatic cell fractions obtained during clinical islet isolation procedures were expanded in culture and subjected to CD81/CD9-based enrichment. Enriched IPC populations were serially passaged to generate monolayer cultures and subsequently formed three-dimensional IPC clusters under high-density conditions. Clusters were isolated and differentiated using ISX9 to generate islet-like organoids. Associated marker expression profiles at each stage are indicated.

**Figure 2.**
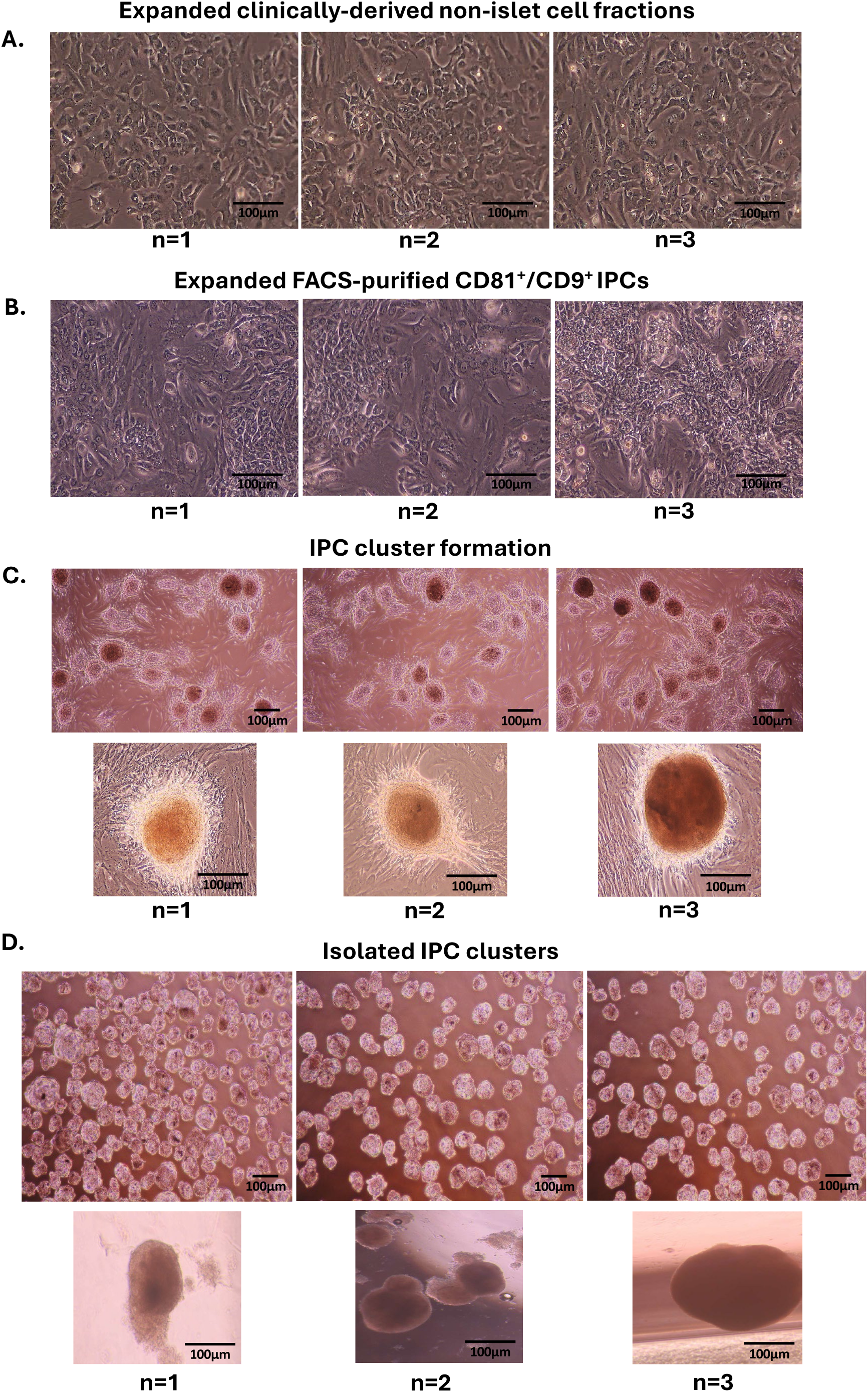
Expansion and cluster formation of clinically derived IPC cultures (A) Representative images of expanded non-endocrine pancreatic cell fractions following 2–3 weeks in culture, demonstrating adherent monolayer morphology. (B) Expanded CD81⁺/CD9⁺ IPCs following enrichment and serial passaging. (C) Formation of three-dimensional IPC clusters from monolayer cultures at high cell density. (D) Isolated IPC clusters following mechanical separation from surrounding monolayer cells. Images are shown from independent donor samples (n=1–3). Scale bars, 100 µm.

### Expanded IPC cultures reproducibly generate three-dimensional clusters from clinically derived samples

To evaluate the morphological characteristics and reproducibility of IPC expansion and cluster formation, we examined cultures derived from multiple independent clinical samples. Expanded non-endocrine pancreatic cell fractions formed adherent monolayers composed of spindle-like and epithelial-like cells following 2–3 weeks in culture (Fig. 2A). Similar morphological features were observed across donors, indicating consistent expansion behavior under these conditions. Following enrichment, expanded CD81^+^/CD9^+^

IPCs maintained comparable monolayer morphology, with dense cellular networks observed across independent samples (Fig. 2B). Upon continued culture at high cell density, these monolayers gave rise to three-dimensional IPC clusters, which appeared as compact, spherical aggregates of varying size (Fig. 2C). Cluster formation was observed reproducibly across all donor samples examined. To further define this population, IPC clusters were mechanically isolated from surrounding monolayer cells. Isolated clusters exhibited a more uniform, compact morphology compared to mixed cultures and were readily distinguishable from adherent monolayer cells (Fig. 2D). These observations demonstrate that clinically derived IPC cultures reproducibly transition from monolayer expansion to three-dimensional cluster formation, and that cluster isolation yields a discrete population suitable for downstream differentiation.

### CD81/CD9-based enrichment defines IPC-associated populations within non-endocrine pancreatic fractions

To characterize the cellular composition of expanded cultures derived from non-endocrine pancreatic cell fractions, we performed flow cytometry analysis prior to downstream expansion and differentiation. Cultures expanded for 2-3 weeks contained a heterogeneous population of cells expressing CD81 and CD9, with subsets co-expressing IPC-associated marker PROCR and islet differentiation factor PDX1, while exhibiting minimal (∼1-2%) expression of endocrine hormones insulin and glucagon (Fig. 3A).

**Figure 3.**
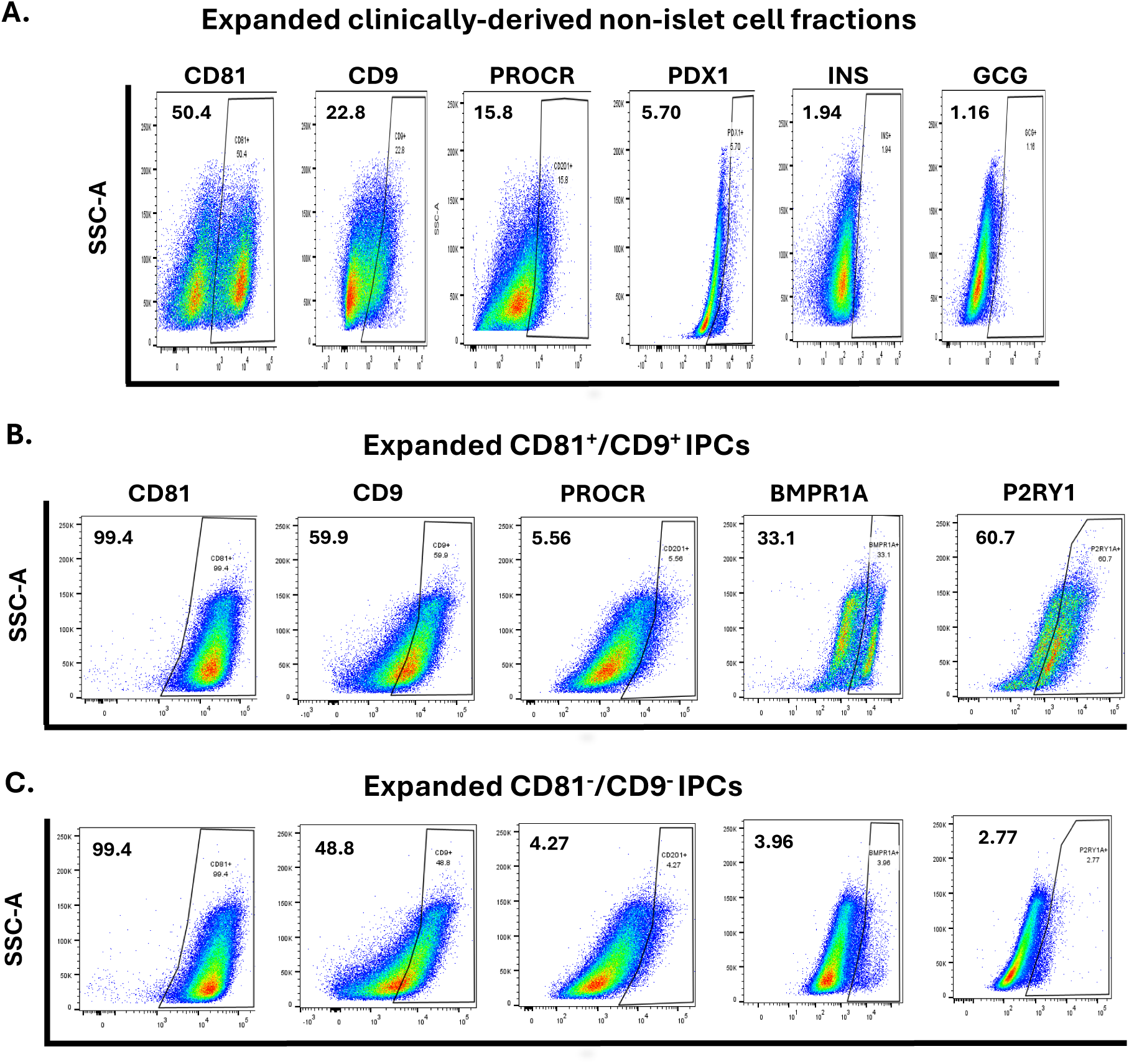
CD81/CD9-based enrichment defines IPC-associated populations (A) Flow cytometry analysis of expanded non-endocrine pancreatic cell fractions showing expression of CD81, CD9, PROCR, PDX1, and low levels of endocrine hormones (INS, GCG). (B) Flow cytometry analysis of expanded CD81⁺/CD9⁺ IPCs demonstrating enrichment of IPC-associated markers, including BMPR1A and P2RY1. (C) Expanded CD81⁻/CD9⁻ populations showing re-expression of CD81 and CD9 but reduced expression of IPC-associated markers. Percentages indicate proportion of positive cells within each population.

FACS-purification of double positive CD81^+^/CD9^+^ IPCs confirmed enriched gene co-expression of CD81 and CD9 with IPC-associated markers PROCR, RGS16, NEUROG3, and PDX1 compared to presorted and double negative CD81^−^/CD9^−^ cell populations (Suppl. Fig 1A). Upon further expansion, CD81^+^/CD9^+^ cells yielded a population that exhibited high expression IPC-associated surface markers BMPR1A (33.1%) and P2RY1 (60.7%) with reduced subset of PROCR (5.56%) (Fig. 3B).

In contrast, CD81/CD9-depleted populations re-expressed CD81(99.4%) and CD9 (48.8%) after 2-3 weeks culture, but exhibited low expression of PROCR (4.27%), BMPR1A (3.96%), and P2RY1 (2.77%) (Fig. 3C). These findings indicate that CD81/CD9-based sorting enables prospective isolation of cell populations enriched for IPC-associated BMPR1A and P2RY1 from heterogeneous non-endocrine pancreatic compartments. They also demonstrate the capacity of purified CD81^−^/CD9^−^ cells to reset proportionate expression of CD81 and CD9 while retaining a depleted population of IPC-associated markers.

### IPC cluster formation is associated with enrichment of progenitor-associated marker expression

To assess molecular differences associated with structural organization, we compared marker expression between IPC monolayer cells and isolated IPC clusters derived from CD81^+^/CD9^+^ and CD81^−^/CD9^−^ populations. Both monolayer and cluster populations retained expression of CD81 and CD9; however, CD81^+^/CD9^+^ IPC clusters demonstrated increased expression of progenitor-associated markers, including BMPR1A (33.2%) and RGS16 (25.2%), relative to monolayer cells and cultures expanded from purified CD81^−^/CD9^−^ populations (Fig. 4A). Gene expression analysis of CD81^+^/CD9^+^ IPC monolayer and clusters confirmed enrichment of progenitor-associated markers BMPR1A, RGS16, NEUROG3, and PDX1 (Suppl. Fig. 1B). This pattern was consistently observed across independent samples, with IPC clusters exhibiting a more pronounced enrichment of these markers compared to corresponding monolayer populations (Fig. 4B). In contrast, monolayer cells showed comparatively lower expression of these features despite maintaining CD81/CD9 positivity. Together, these data indicate that cluster formation is associated with a distinct molecular profile, supporting the use of CD81^+^/CD9^+^-enriched IPC clusters as a defined substrate for downstream differentiation analyses.

**Figure 4.**
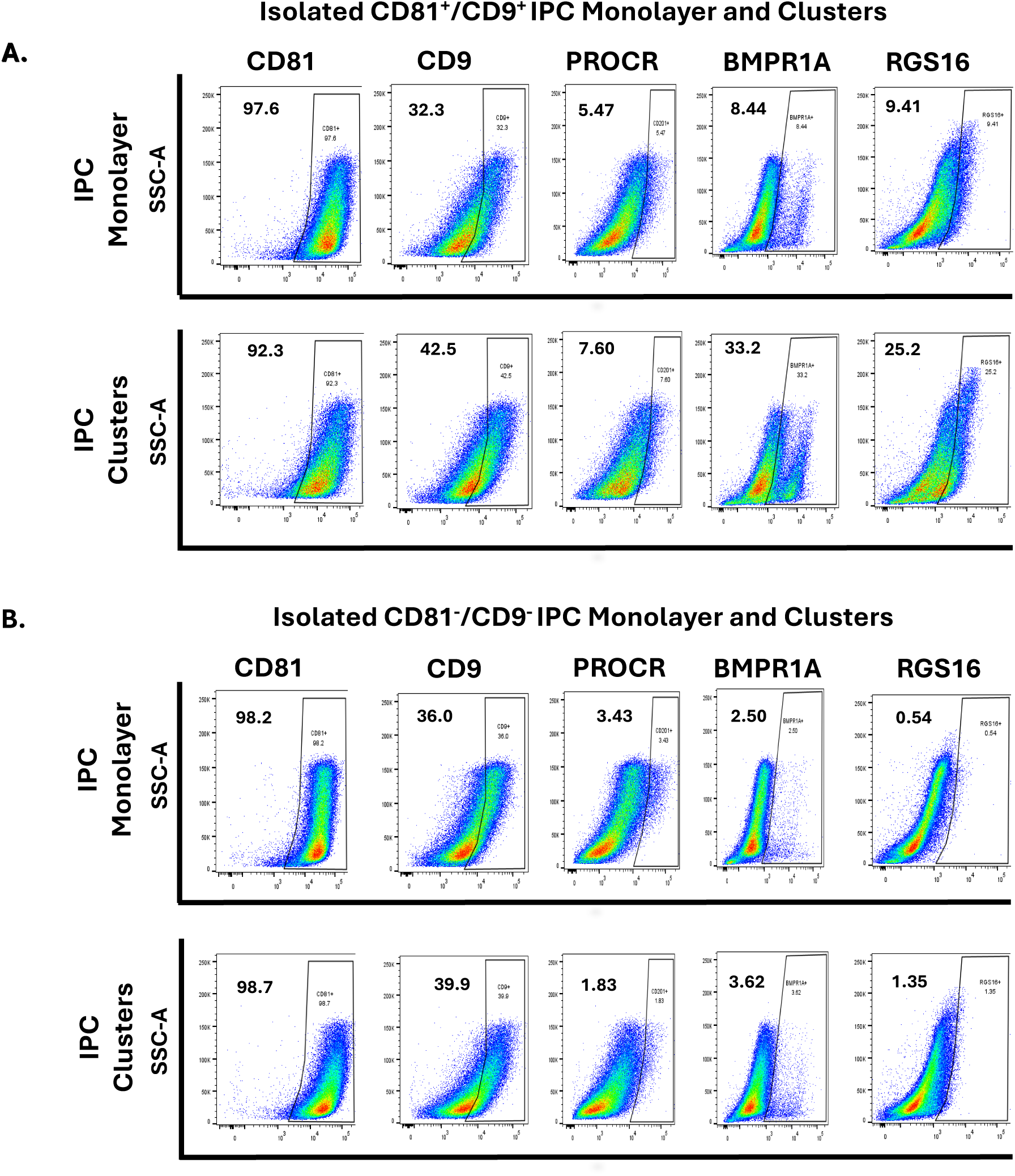
IPC cluster formation is associated with enrichment of progenitor-associated markers (A) Flow cytometry comparison of CD81⁺/CD9⁺ IPC monolayer cells and isolated IPC clusters, demonstrating increased expression of BMPR1A and RGS16 in clusters. (B) Comparison of CD81⁻/CD9⁻ monolayer and cluster populations, showing reduced expression of progenitor-associated markers relative to CD81⁺/CD9⁺ populations. Percentages indicate proportion of positive cells within each population.

### Transcriptomic profiling defines IPC-associated states and progression toward endocrine cell identities

To further define the molecular identity of IPC populations and their progression toward endocrine differentiation, we performed single-cell RNA sequencing across sequential stages of the workflow, including IPC monolayer cultures, IPC clusters, and ISX9-treated organoids. Unsupervised clustering identified distinct cell populations corresponding to IPCs, progenitor-like IPCs, poly-endocrine (Poly-Endo), and lineage-resolved endocrine cell types, including beta-, alpha-, and delta-like populations, alongside smaller non-endocrine compartments such as ductal-like, acinar, endothelial, and immune-like cells (Fig. 5C). IPC populations formed a broad expanded compartment, while progenitor-like IPCs represented a transcriptionally distinct subset within this space. IPC-associated populations were characterized by expression of genes linked to undifferentiated pancreatic states, including GREM1, FST, PTX3, and CEMIP, which were broadly enriched across IPC compartments and reduced following endocrine differentiation (Fig. 5A,B).

**Figure 5.**
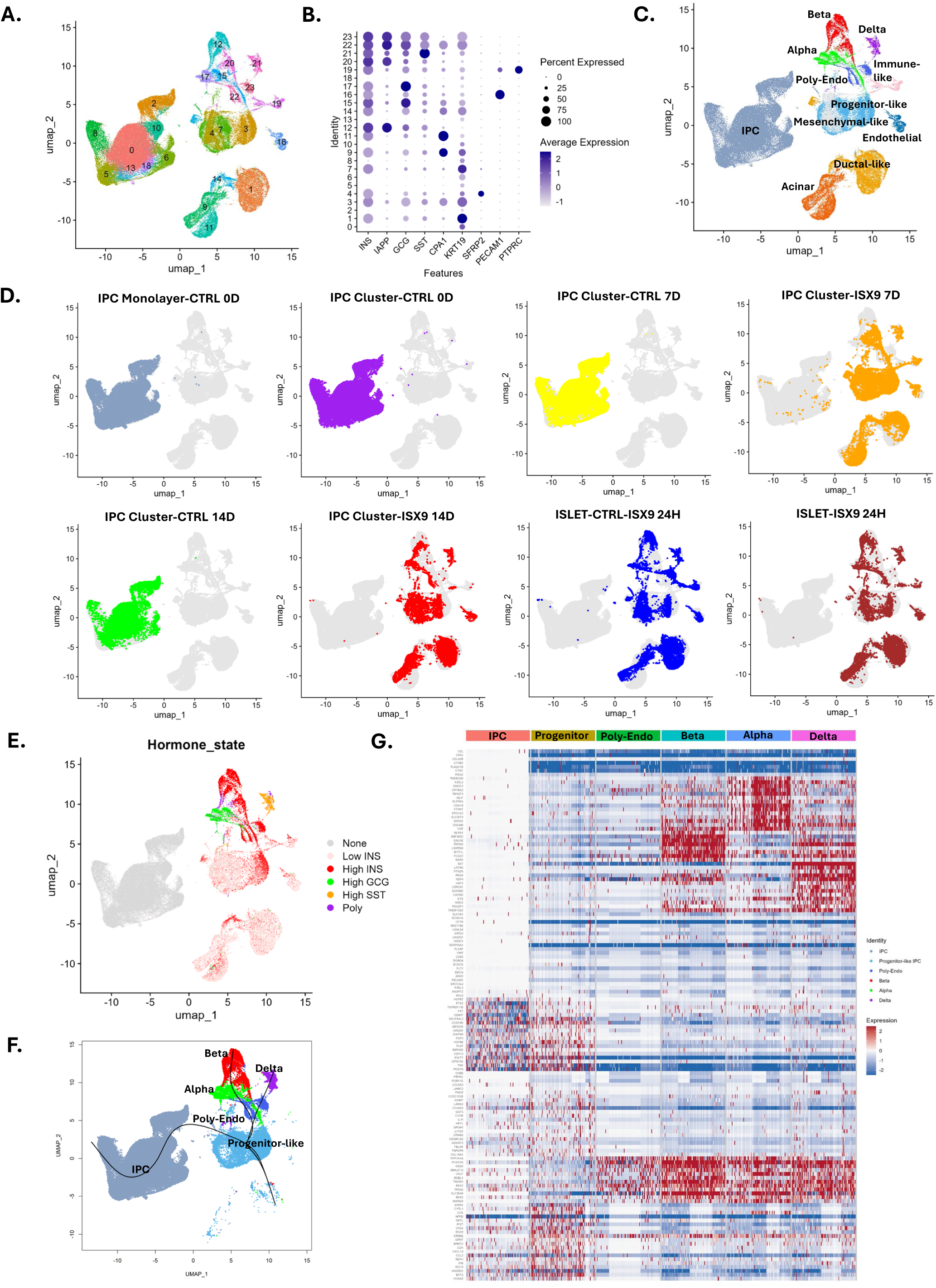
Single-cell transcriptomic profiling defines IPC-associated states and endocrine cell populations (A) Dot plot showing expression of selected genes across experimental conditions, including IPC monolayer, IPC clusters, ISX9-treated clusters, and islet-like populations. (B) Feature plots illustrating expression of IPC-associated genes (CD81, CD9, BMPR1A) and IPC-associated markers (GREM1, FST, PTX3, CEMIP), as well as endocrine markers (INS, GCG, IAPP, SST). (C) UMAP visualization of all cells colored by annotated cell type, including IPC, progenitor-like IPC, poly-endocrine, beta-, alpha-, and delta-like populations, and non-endocrine cell types. (D) UMAP visualization colored by experimental condition, showing progression from IPC monolayer and control clusters to ISX9-treated clusters and islet-like populations. (E) Hormone expression overlay identifying polyhormonal (Poly-Endo) and monohormonal endocrine cell populations. (F) Projection of lineage relationships indicating a branching structure from Poly-Endo populations toward beta-and alpha-like cell types. (G) Heatmap showing coordinated expression of IPC-associated and endocrine gene programs across annotated cell types.

Mapping of cells by experimental condition demonstrated a clear progression across the workflow. Early-stage IPC monolayer and control cluster populations localized predominantly within IPC and progenitor-like regions of the UMAP, whereas cells from later differentiation stages, particularly IPC Cluster-ISX9 14D and ISLET-ISX9 24H, localized within endocrine regions and co-clustered with beta-, alpha-, delta-, and poly-endocrine populations (Fig. 5D).

Analysis of hormone expression patterns further supported this transition. A polyhormonal intermediate population (Poly-Endo) was identified, characterized by co-expression of endocrine hormones and positioned between IPC/progenitor-like compartments and lineage-resolved endocrine cell types (Fig. 5E). Monohormonal insulin-and glucagon-expressing populations were predominantly observed in later differentiation stages, consistent with progression toward more defined endocrine identities.

Projection of lineage relationships suggested a branching structure in which Poly-Endo populations are positioned upstream of beta-and alpha-like populations (Fig. 5F). While these relationships are not intended to establish definitive lineage trajectories, they are consistent with transitional endocrine states emerging from IPC-derived populations.

Consistent with these observations, heatmap analysis demonstrated coordinated expression of progenitor-associated gene programs in IPC and progenitor-like populations, including GREM1, FST, PTX3, and CEMIP, and activation of endocrine transcriptional and functional gene programs in beta-, alpha-, and delta-like clusters, including INS, GCG, IAPP, and SST (Fig. 5G). Global transcriptional patterns across all conditions further support these transitions, as shown by expanded heatmap analysis (Suppl. Fig. 2A,B).

Together, these data demonstrate that IPC populations derived from non-endocrine pancreatic cell fractions occupy transcriptionally defined intermediate states that are preserved during expansion and can be directed toward endocrine cell identities, resulting in organoid populations that exhibit islet-like transcriptomic profiles.

### Gene program analysis and feature mapping distinguish IPC-associated and endocrine transcriptional states

To further define transcriptional programs associated with IPC and endocrine cell states, we examined expression of grouped gene sets and their spatial distribution across the dataset.

Across experimental conditions, distinct gene expression programs differentiated IPC-associated and endocrine populations (Fig. 6A). IPC monolayer and cluster populations were enriched for markers associated with undifferentiated states, including CD81, CD9, and BMPR1A, as well as IPC-associated genes GREM1, FST, PTX3, and CEMIP. In contrast, endocrine populations were defined by expression of hormone genes, including INS, GCG, IAPP, and SST. Expanded analysis across all clusters and annotated cell types further supports these patterns (Suppl. Fig. 3A).

**Figure 6.**
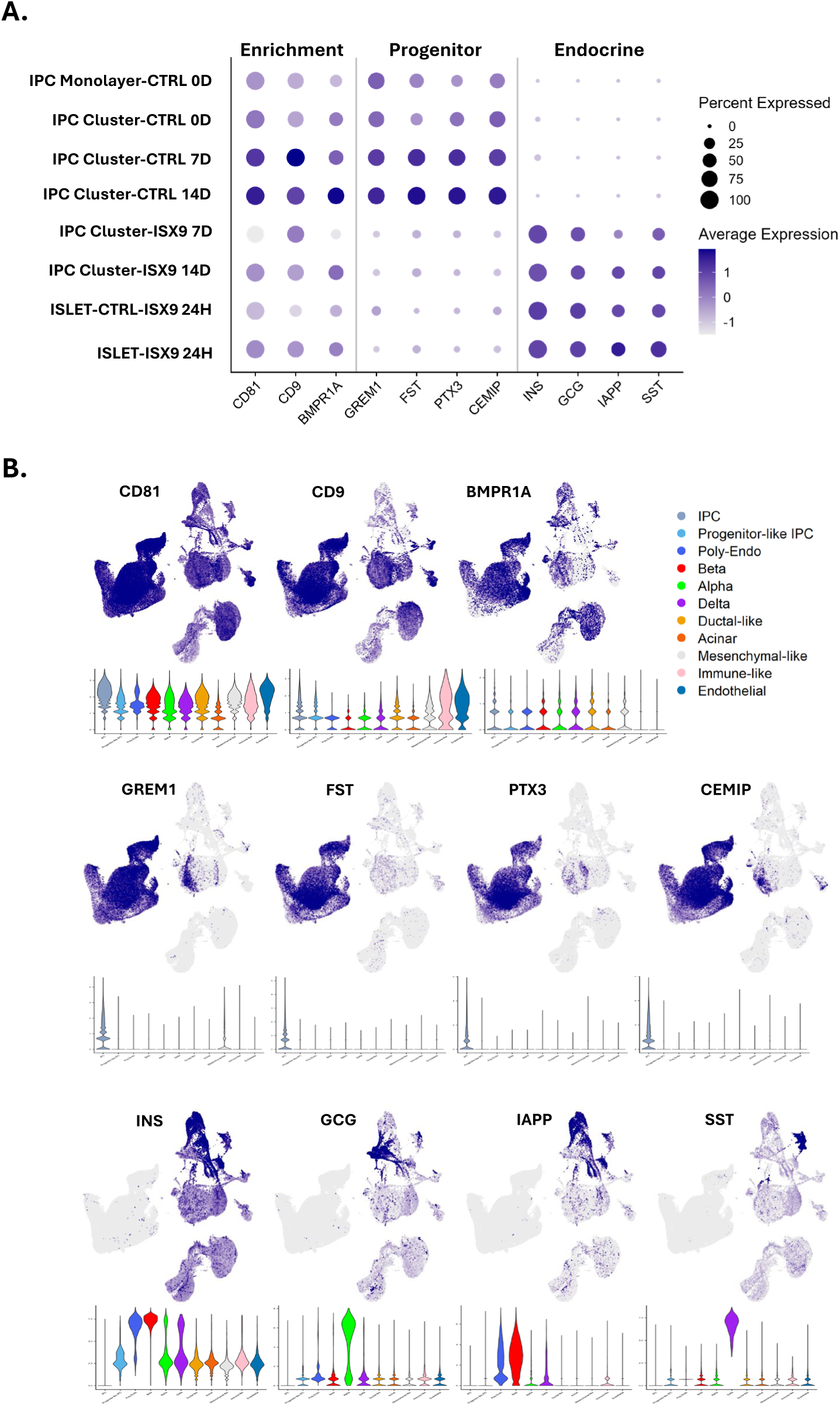
Gene program analysis and feature mapping distinguish IPC-associated and endocrine states (A) Dot plot showing expression of enrichment-associated, IPC-associated, and endocrine gene sets across experimental conditions. (B) Feature plots illustrating spatial distribution of IPC-associated markers (CD81, CD9, BMPR1A), IPC-associated genes (GREM1, FST, PTX3, CEMIP), and endocrine markers (INS, GCG, IAPP, SST) across the UMAP. Data demonstrate distinct transcriptional programs associated with IPC and endocrine cell populations.

Feature mapping across the UMAP further reinforced these observations (Fig. 6B). IPC-associated markers, including CD81, CD9, and BMPR1A, localized predominantly to IPC and progenitor-like regions, while IPC-associated genes GREM1, FST, PTX3, and CEMIP were broadly enriched across these compartments. In contrast, endocrine markers (INS, GCG, IAPP, and SST) were restricted to discrete clusters corresponding to lineage-resolved beta-, alpha-, and delta-like populations.

Together, these analyses demonstrate that IPC populations exhibit coordinated transcriptional programs that distinguish IPC-associated and endocrine states, and that these programs shift in a stage-dependent manner during differentiation. These patterns are consistent with the emergence of islet-like endocrine cell populations following directed differentiation of IPC-derived clusters.

### ISX9 induces stage-dependent expression of endocrine transcription factors and hormone-processing genes in IPC-derived clusters

To evaluate endocrine differentiation at the molecular level, we assessed gene expression of key islet-associated transcription factors and functional markers in IPC-derived clusters following ISX9 treatment across multiple time points. Expression of canonical beta cell transcription factors, including PDX1, NKX2.2, MAFA, and NKX6.1, was induced within 8 days following ISX9 treatment compared to controls. Notably, increases in PDX1 and MAFA expression were observed at later time points (Fig. 7A). Expression of additional regulatory factors NKX2.2 and NKX6.1 showed more modest or variable changes over time.

**Figure 7.**
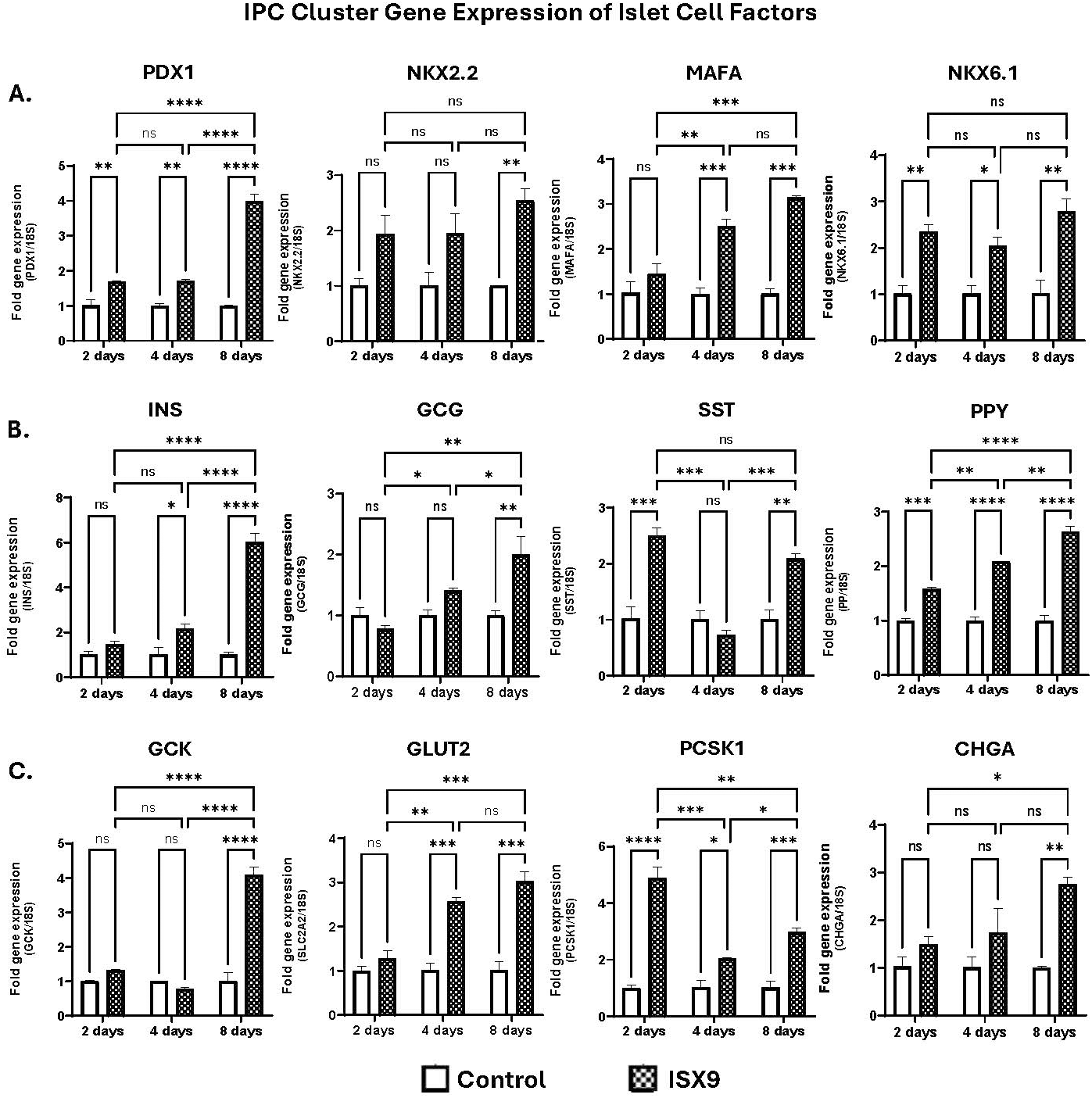
ISX9 induces endocrine transcription factor expression and functional gene programs in IPC-derived clusters (A) Expression of endocrine transcription factors (PDX1, NKX2.2, MAFA, NKX6.1) in IPC clusters following ISX9 treatment at 2, 4, and 8 days. (B) Expression of hormone genes (INS, GCG, SST, PPY) following ISX9 treatment across time points. (C) Expression of genes associated with glucose sensing and hormone processing (GCK, SLC2A2/GLUT2, PCSK1, CHGA). Data are presented as fold change relative to control. Statistical significance indicated as shown.

Consistent with induction of endocrine identity, ISX9-treated clusters exhibited increased expression of hormone genes, including INS and GCG, with progressive increases observed from 2 to 8 days (Fig. 7B). Expression of additional endocrine markers, including PPY, also increased, whereas SST expression showed more limited or variable induction.

Analysis of genes associated with glucose sensing and hormone processing further supported functional maturation. Expression of GCK, GLUT2, and PCSK1 was increased in ISX9-treated clusters relative to controls, indicating acquisition of key components of glucose responsiveness and prohormone processing pathways (Fig. 7C). Expression of CHGA was also elevated, consistent with development of endocrine secretory machinery.

### IPC-derived islet organoids exhibit increased insulin-and glucagon-expressing cell populations following differentiation

To assess the cellular composition of differentiated organoids, we performed flow cytometry and immunofluorescence analysis of hormone-expressing cells across differentiation time points. ISX9-treated islet organoids demonstrated a marked increase in the proportion of INS⁺ cells compared to control conditions, with progressive enrichment observed from 2 to 8 days (Fig. 8A). Similarly, GCG⁺ cell populations were increased following ISX9 treatment, indicating generation of multiple endocrine cell types.

**Figure 8.**
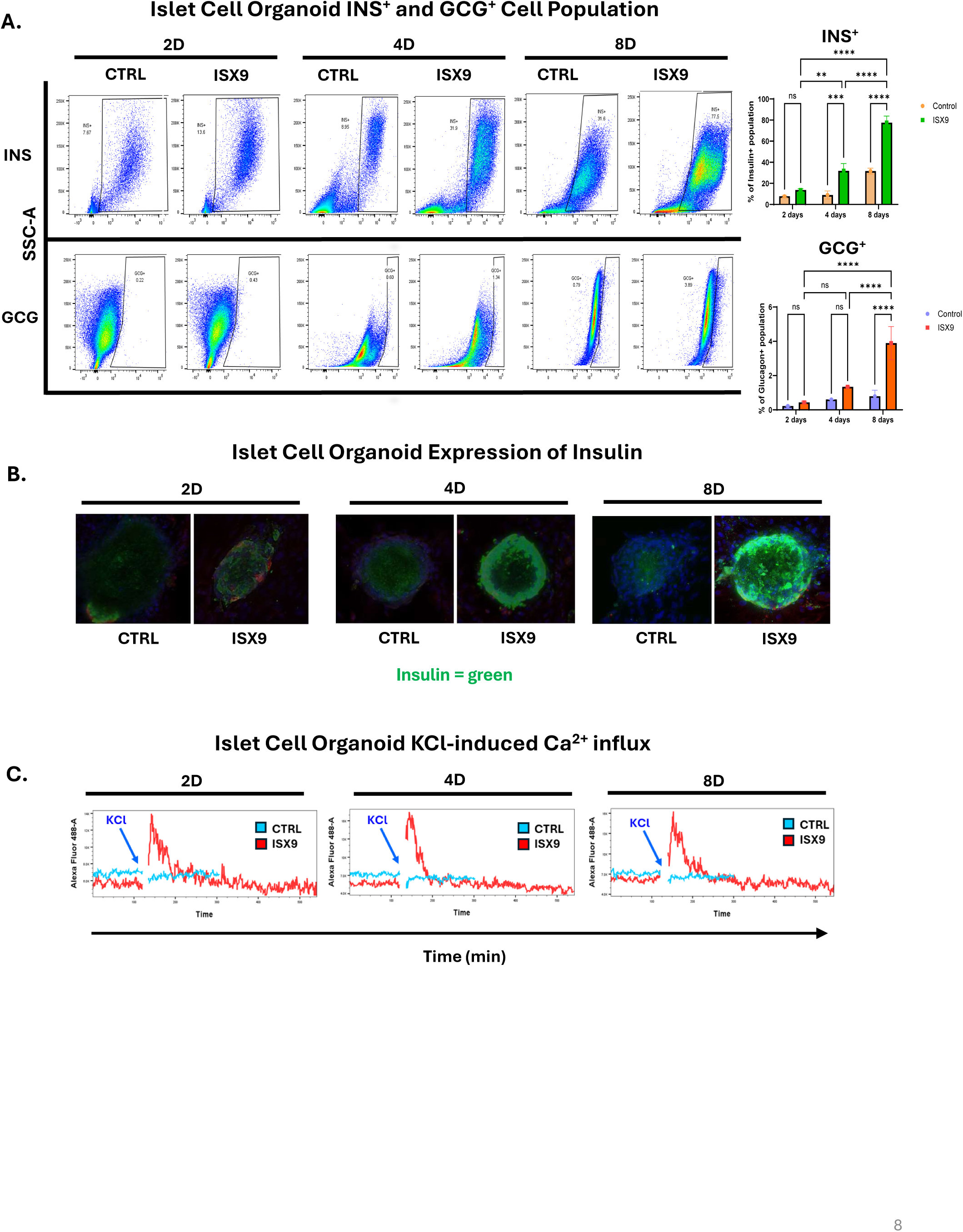
IPC-derived organoids exhibit endocrine cell populations and functional calcium responses (A) Flow cytometry analysis of INS⁺ and GCG⁺ cell populations in control and ISX9-treated organoids across time points. (B) Immunofluorescence images of insulin expression (green) in IPC-derived organoids under control and ISX9 conditions. (C) Intracellular calcium dynamics in response to KCl stimulation, demonstrating depolarization-induced Ca²⁺ influx in ISX9-treated organoids.

Immunofluorescence analysis confirmed these findings, with ISX9-treated islet organoids exhibiting increased insulin-positive staining and more defined three-dimensional structures compared to control conditions (Fig. 8B). These observations are consistent with differentiation of IPC-derived clusters into hormone-expressing endocrine-like organoids.

### Differentiated IPC-derived organoids exhibit stimulus-responsive calcium signaling

To assess functional responsiveness, we evaluated calcium dynamics in IPC-derived islet organoids following depolarization. ISX9-treated organoids exhibited robust KCl-induced Ca²⁺ influx, with clear increases in intracellular calcium levels following stimulation across differentiation time points (Fig. 8C). In contrast, control IPC clusters were unresponsive to KCl stimulation, indicating a lack of depolarization-induced calcium influx. These findings indicate that IPC-derived islet organoids acquire functional ion channel activity associated with endocrine cell excitability following differentiation.

### IPC-derived islet organoids exhibit glucose-responsive hormone secretion

To further assess functional maturation, we evaluated hormone secretion in response to glucose stimulation. ISX9-treated islet organoids demonstrated increased insulin secretion under high-glucose conditions compared to low-glucose conditions, consistent with acquisition of glucose responsiveness (Fig. 9). In parallel, glucagon secretion was elevated under low-glucose conditions, reflecting appropriate counter-regulatory hormone release. Together, these findings demonstrate that IPC-derived organoids acquire key functional properties of endocrine cells, including stimulus-responsive calcium signaling and glucose-regulated hormone secretion.

**Figure 9.**
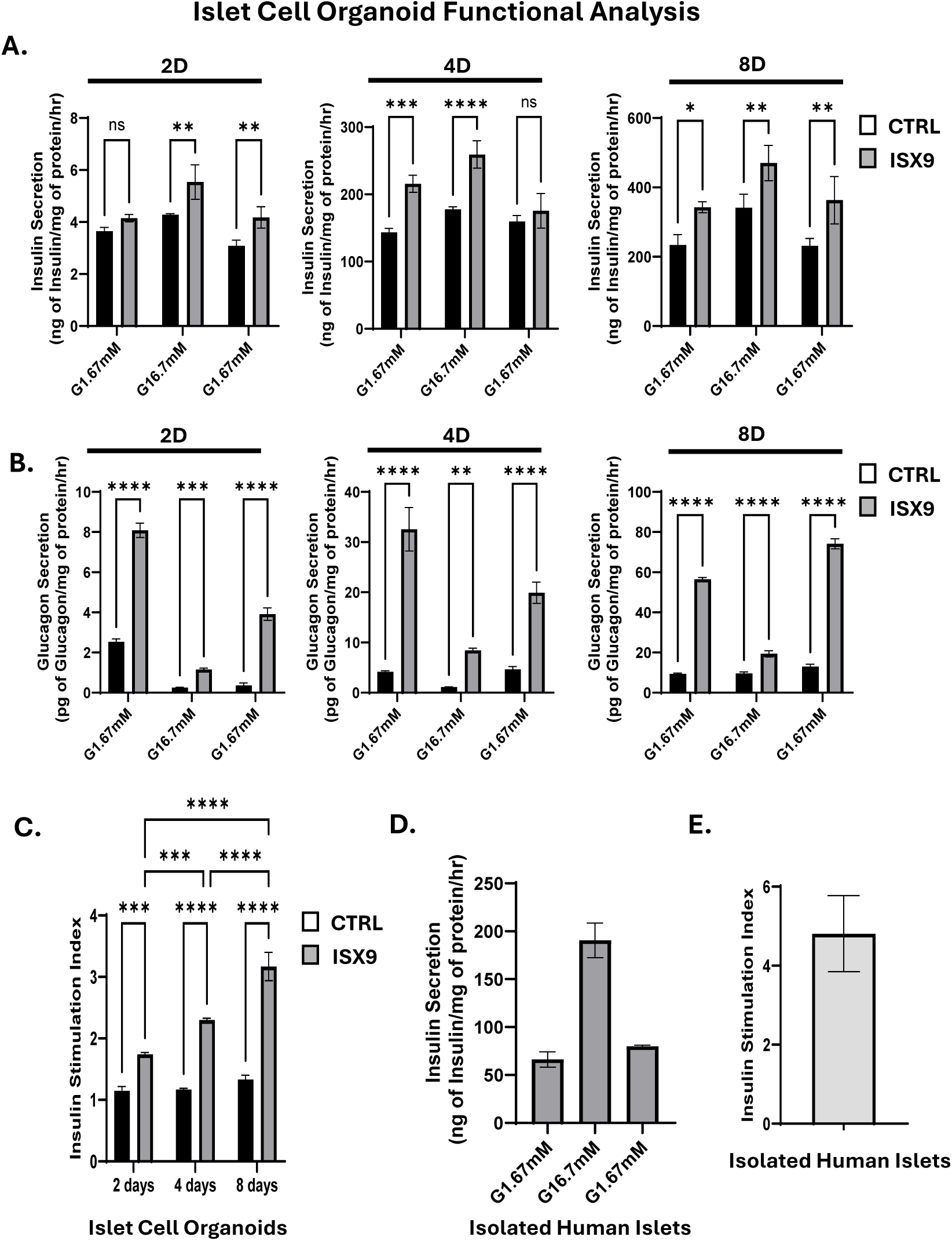
IPC-derived organoids exhibit glucose-responsive hormone secretion (A–C) Insulin secretion in response to low (1.67 mM) and high (16.7 mM) glucose conditions across differentiation time points. (D–E) Glucagon secretion under low-and high-glucose conditions. (F) Comparison of insulin secretion from IPC-derived organoids and isolated human islets. (G) Insulin stimulation index demonstrating glucose responsiveness in control and ISX9-treated organoids. Data are normalized to protein content. Statistical significance indicated as shown.

## DISCUSSION

In this study, we establish a clinically adaptable platform for the expansion, enrichment, and directed differentiation of adult pancreas-derived progenitor populations obtained from pancreatic tissue processed during clinical islet isolation. We demonstrate that non-endocrine fractions generated during islet processing contain cells capable of expansion into IPC populations, which can be enriched for islet differentiation potential using CD81/CD9-based sorting and subsequently differentiated into islet-like organoids exhibiting endocrine transcriptional programs and functional properties. These findings extend our prior work by translating an experimentally defined progenitor platform into a workflow compatible with clinically derived tissue sources.

A central advance of this study is the implementation of a prospective enrichment strategy. While prior work identified CD81 and CD9 as highly expressed in expanded IPC populations, here we demonstrate that CD81/CD9-based sorting reproducibly enriches a population with enhanced capacity for cluster formation and downstream endocrine differentiation. Importantly, enriched IPC populations exhibit increased expression of markers associated with progenitor competency, including BMPR1A and RGS16, and demonstrate improved efficiency in generating structured IPC clusters. In contrast, CD81/CD9-depleted populations, despite re-expression of these markers during expansion, do not recover comparable progenitor-associated features, suggesting that early enrichment enhances differentiation potential rather than simply selecting for culture-adapted cells.

Single-cell transcriptomic profiling further defines the cellular architecture underlying IPC expansion and differentiation. IPC populations were characterized by expression of genes associated with undifferentiated or plastic cellular states, including GREM1, FST, PTX3, and CEMIP, which were progressively reduced following differentiation. Upon ISX9 treatment, IPC-derived clusters transitioned toward endocrine populations exhibiting transcriptional profiles consistent with beta-, alpha-, and delta-like cell identities. Importantly, differentiated IPC clusters localized within endocrine regions of transcriptomic space and aligned with human islet reference signatures, supporting acquisition of islet-like identity at the molecular level. While these analyses were performed on a limited number of donors and should be interpreted as representative, they provide high-resolution evidence of coordinated endocrine program activation.

Functional analyses demonstrate that IPC-derived organoids acquire key features of endocrine physiology. Differentiated organoids exhibited depolarization-induced calcium influx and glucose-regulated insulin and glucagon secretion, indicating acquisition of stimulus-responsive signaling and hormone release mechanisms. Although secretion magnitude and stimulation indices differed from primary human islets, the presence of dynamic glucose responsiveness supports the conclusion that IPC-derived organoids achieve a functionally relevant endocrine state within a relatively short differentiation window.

The broader implications of this platform extend beyond the specific clinical context in which these tissues were obtained. While non-islet fractions generated during islet isolation provide a practical and clinically relevant source of starting material, these studies establish a proof-of-concept that adult human pancreatic tissue contains cells capable of expansion and directed differentiation into endocrine-like organoids. This raises the possibility that small amounts of patient-derived pancreatic tissue could be expanded ex vivo to generate functional endocrine cells, providing a potential framework for autologous cell replacement strategies in diabetes, including type 1 diabetes.

Several limitations should be considered. Surface marker expression is influenced by ex vivo culture conditions, and CD81/CD9 should be interpreted as an operational enrichment strategy rather than definitive identification of an endogenous progenitor lineage.

Transcriptomic analyses were performed on a limited number of donors, and further studies will be required to assess inter-donor variability and reproducibility. In addition, while in vitro functional assays demonstrate acquisition of key endocrine properties, the long-term stability, maturation, and in vivo performance of these organoids remain to be fully defined.

In summary, we present a clinically adaptable workflow that converts pancreatic tissue fractions generated during islet isolation into functional islet-like organoids. CD81/CD9-based enrichment enhances recovery of endocrine-competent IPC populations and supports reproducible differentiation into organoids exhibiting coordinated transcriptional and functional features of islet cells. This strategy provides a foundation for future efforts to expand autologous pancreatic cell populations for therapeutic applications in diabetes.\

---

**Supplementary Figure 1.**
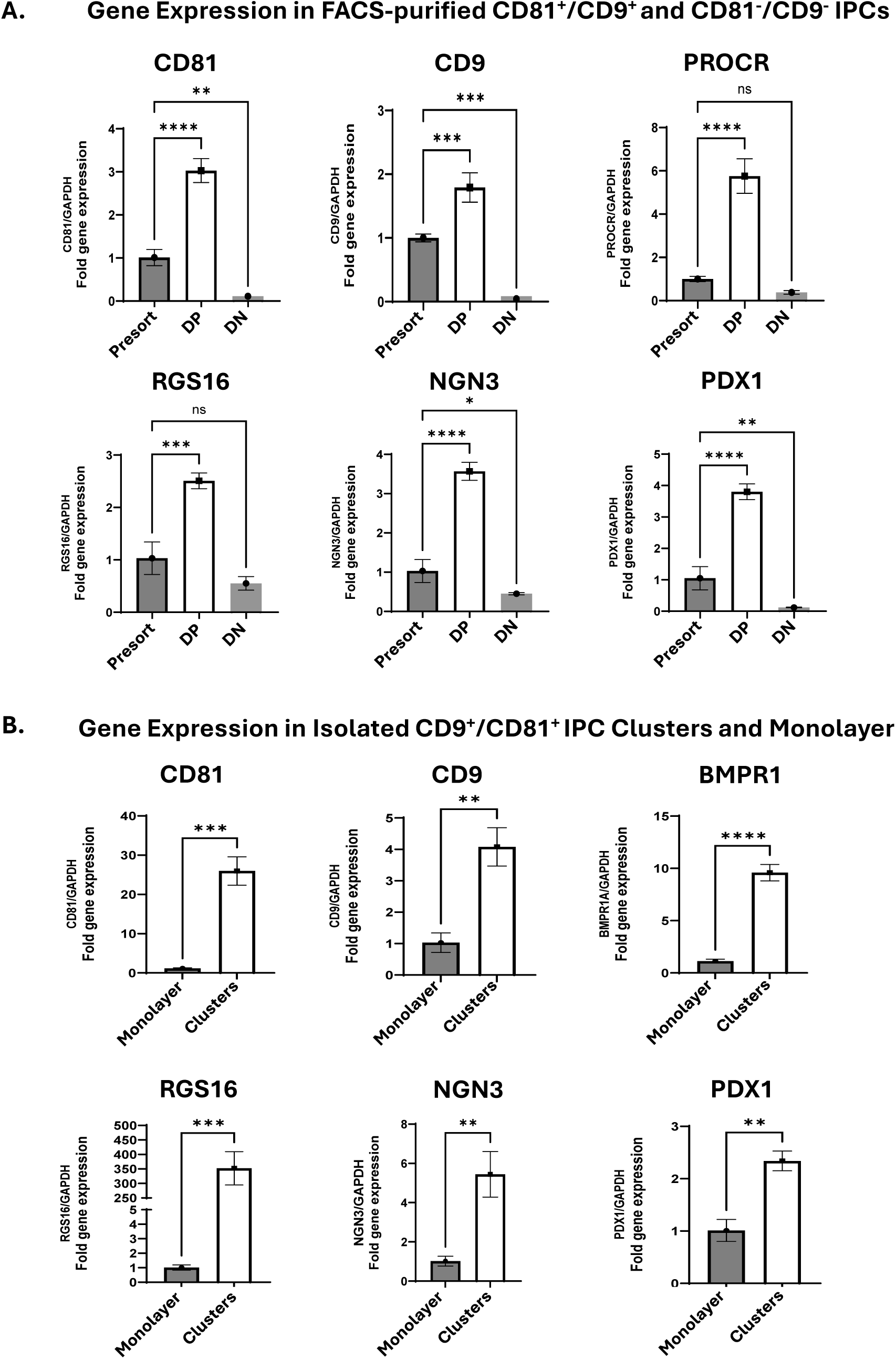
Gene expression analysis of enriched IPC populations (A) Gene expression comparison of unsorted, CD81⁺/CD9⁺ (DP), and CD81⁻/CD9⁻ (DN) populations. (B) Gene expression comparison of IPC monolayer and cluster populations. Expression of CD81, CD9, PROCR, BMPR1A, RGS16, NGN3, and PDX1 is shown.

**Supplementary Figure 2.**
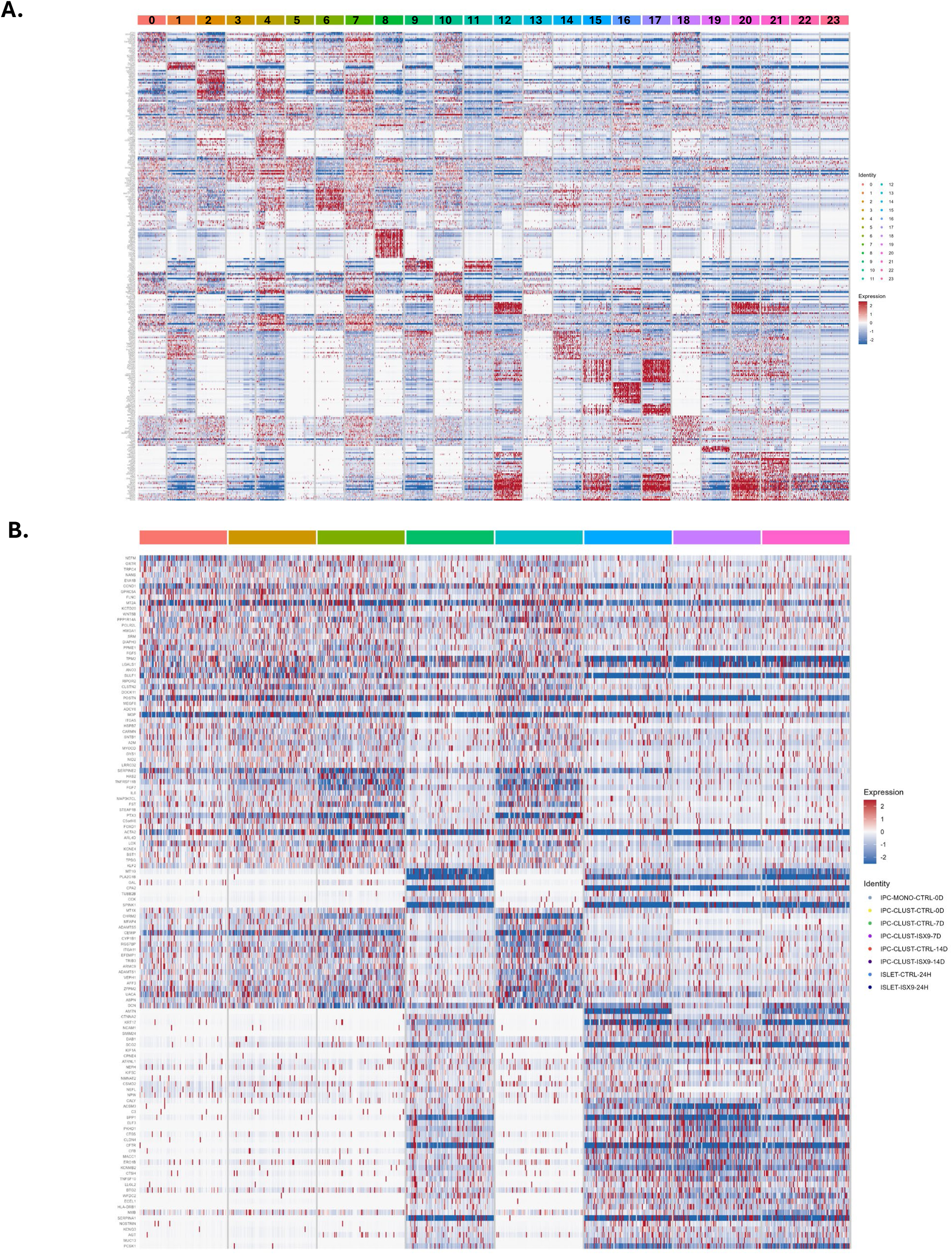
Global transcriptional heatmap across experimental conditions Heatmaps showing global gene expression patterns across (A) all clusters 1-23 and annotated cell types, supporting coordinated transitions between IPC-associated and endocrine gene programs.

**Supplementary Figure 3.**
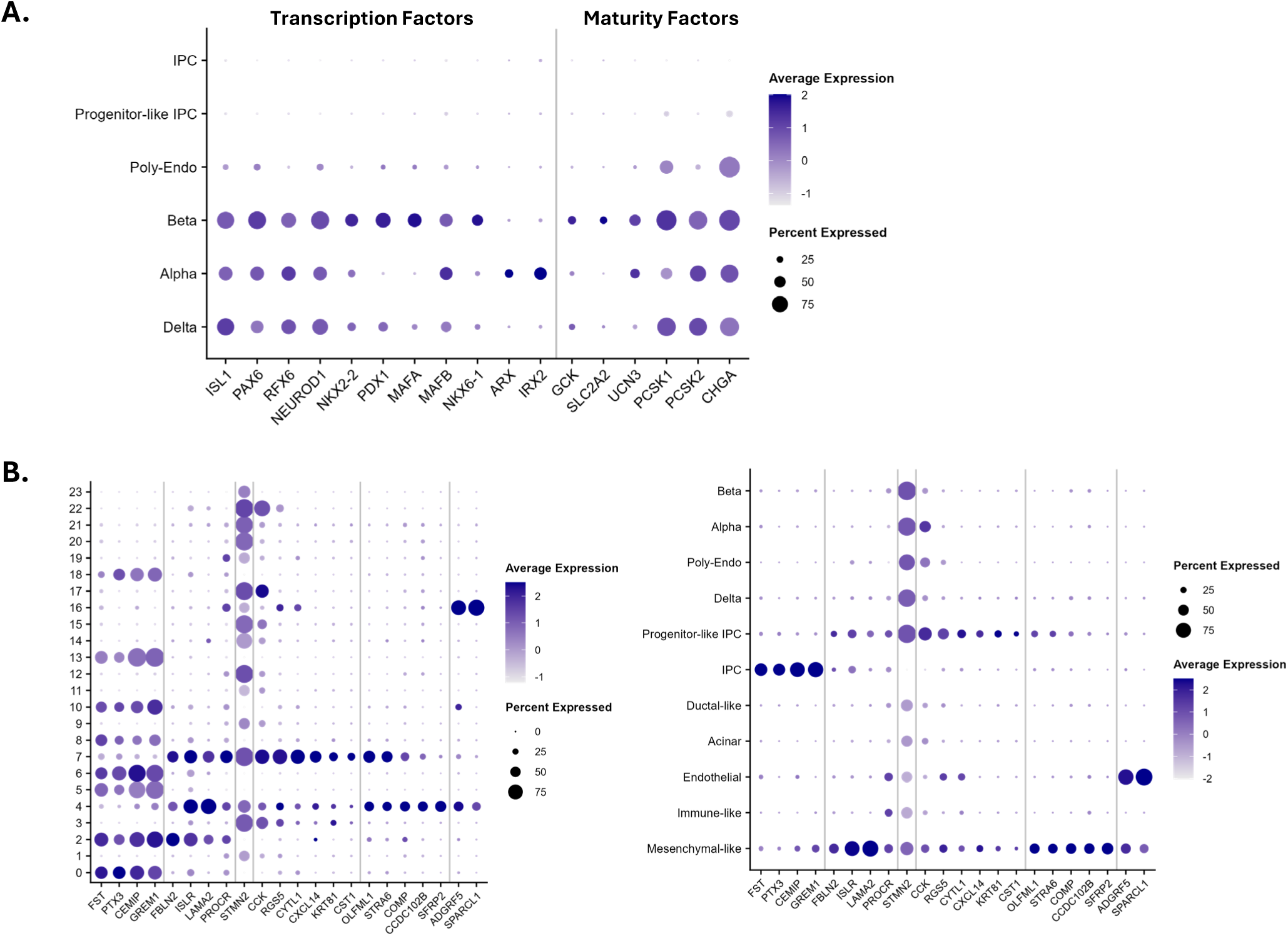
Expanded gene program analysis and endocrine transcription factor expression (B) Dot plot showing expression of gene programs across all clusters and annotated cell types. (C) Dot plot of islet-associated transcription factors and maturation-related genes across annotated cell types.

## REFERENCES

1. A. M. Shapiro et al., International trial of the Edmonton protocol for islet transplantation. N Engl J Med 355, 1318–1330 (2006).

2. F. B. Barton et al., Improvement in outcomes of clinical islet transplantation: 1999-2010. Diabetes Care 35, 1436–1445 (2012).

3. D. E. Sutherland et al., Islet autotransplant outcomes after total pancreatectomy: a contrast to islet allograft outcomes. Transplantation 86, 1799–1802 (2008).

4. Y. Wang et al., The future of islet transplantation beyond the BLA approval: challenges and opportunities. Front Transplant 4, 1522409 (2025).

5. A. Rezania et al., Reversal of diabetes with insulin-producing cells derived in vitro from human pluripotent stem cells. Nat Biotechnol 32, 1121–1133 (2014).

6. F. W. Pagliuca et al., Generation of functional human pancreatic beta cells in vitro. Cell 159, 428–439 (2014).

7. D. Balboa et al., Functional, metabolic and transcriptional maturation of human pancreatic islets derived from stem cells. Nat Biotechnol 40, 1042–1055 (2022).

8. T. W. Reichman et al., Stem Cell-Derived, Fully Differentiated Islets for Type 1 Diabetes. N Engl J Med 393, 858–868 (2025).

9. C. Talchai, S. Xuan, H. V. Lin, L. Sussel, D. Accili, Pancreatic beta cell dedifferentiation as a mechanism of diabetic beta cell failure. Cell 150, 1223–1234 (2012).

10. A. Migliorini, E. Bader, H. Lickert, Islet cell plasticity and regeneration. Mol Metab 3, 268–274 (2014).

11. M. M. F. Qadir et al., P2RY1/ALK3-Expressing Cells within the Adult Human Exocrine Pancreas Are BMP-7 Expandable and Exhibit Progenitor-like Characteristics. Cell Rep 22, 2408–2420 (2018).

12. M. M. F. Qadir et al., Single-cell resolution analysis of the human pancreatic ductal progenitor cell niche. Proc Natl Acad Sci U S A 117, 10876–10887 (2020).

13. M. M. F. Qadir et al., Long-term culture of human pancreatic slices as a model to study real-time islet regeneration. Nat Commun 11, 3265 (2020).

14. D. Wang et al., Long-Term Expansion of Pancreatic Islet Organoids from Resident Procr(+) Progenitors. Cell 180, 1198–1211 e1119 (2020).

15. C. Salinno et al., CD81 marks immature and dedifferentiated pancreatic beta-cells. Mol Metab 49, 101188 (2021).

16. C. M. Darden et al., Calcineurin/NFATc2 and PI3K/AKT signaling maintains beta-cell identity and function during metabolic and inflammatory stress. iScience 25, 104125 (2022).

17. C. M. Darden et al., Mechanisms inducing differentiation of adult islet progenitor-like cells into functional islet-like organoids. Front Transplant 5, 1740314 (2026).

